# Meta-omics analyses of dual substrate enrichment culturing of nitrous oxide respiring bacteria suggest that attachment and complex polysaccharide utilisation contributed to the ability of *Cloacibacterium* strains to reach dominance

**DOI:** 10.1101/2023.06.04.543644

**Authors:** Silas H.W. Vick, Kjell Rune Jonassen, Magnus Ø. Arntzen, Pawel Lycus, Lars R Bakken

## Abstract

Bioengineering soil metabolism by inoculation is an emerging approach to enhance plant growth and strengthen specific functions such as N_2_O reduction in order to reduce climate forcing. The use of organic wastes as substrates and carriers of microbial biomass has proven to be a viable approach to improving effectiveness and economic viability. A key factor in the success of this approach lies in selection of microbes capable of growth and survival in both organic wastes as well as soils, and which are tolerant of the rapid environmental fluctuations such fertilisations involve. A dual substrate, N_2_O-enrichment experiment, switching between soil and organic waste as substrates has yielded *Cloacibacterium* isolates which grow well in organic wastes and retain significant N_2_O reduction capacity when applied to soils. However, an understanding of the genetic and phenotypic characteristics utilised by these enrichment winners to dominate under such conditions remains unexplored. Here we have performed a multi-omics examination of the enrichment cultures, using both metagenomics and metaproteomics to probe the genetic basis and expressed proteins which may contribute to the success of *Cloacibacterium* in the enrichments, and their survival in soil. These omics results show an increase in complex carbohydrate metabolism, chemotaxis and motility genes throughout the enrichment as well as the expression of gliding motility proteins and polysaccharide utilization loci proteins by *Cloacibacterium* organisms. Taken together this suggests that attachment and complex polysaccharide utilisation may be key processes allowing *Cloacibacterium* to tolerate the stresses of a changing environment during transfers between digestate and soil.

## Introduction

N_2_O is a powerful and long-lived global warming and ozone depleting gas with a global warming potential 300 times that of CO_2_ (Solomon et al. 2007; Wuebbles 2009). Atmospheric N_2_O emissions are increasing due to human activities, primarily the growing use of nitrogenous fertilisers in agriculture (Davidson 2009; Thompson et al. 2019), where farmed soils account for 52% of all anthropogenic N_2_O emissions (Tian et al. 2020). These N_2_O emissions contribute to a third of the total climate forcing from agriculture (Robertson 2014) and in contrast to other global warming gas emissions few mitigation options are currently available beyond a more constrained use of fertilisers (Winiwarter et al. 2018).

Production of N_2_O from nitrogenous fertilisers in soils is primarily through the microbial processes of nitrification and denitrification, where denitrification is thought to be the dominant source under most agricultural conditions (Butterbach-Bahl et al. 2013a). Denitrification is carried out by diverse facultatively anaerobic microorganisms in response to oxygen depletion (Butterbach-Bahl et al. 2013b), and refers to anaerobic respiration utilising NO_3_ ^-^, NO_2_, NO and N_2_O as terminal electron acceptors in a stepwise manner, resulting in final conversion to N_2_. These reactions are catalysed by a series of reductase enzymes: the NO_3_ ^-^ reductases NapA and NarG, the NO_2_ ^-^ reductases NirK and NirS, the NO reductases cNor and qNor and the N_2_O reductase NosZ, which is divided into clade I or II (Hallin et al. 2018). Denitrifying organisms can carry a full suite of denitrification reductases or a truncated subset (Shapleigh 2013; Lycus et al. 2017). The abundance and activity of truncated denitrifiers, along with their regulatory phenotypes, can influence the ratio of gaseous N_2_O and N_2_ escaping the soil (Philippot et al 2011).

Recently, it has been demonstrated that soil N_2_O emissions can be altered through heavy inoculation with truncated denitrifiers carrying only the N_2_O reductase gene *nosZ*, termed N_2_O reducing bacteria (NRB) (Philippot et al. 2011; Domeignoz-Horta et al. 2016; Gao et al. 2017). While such heavy inoculation with NRB grown in fermenters would be complicated and expensive, a feasible alternative would be to grow such organisms in organic wastes, which are destined to be used as biofertilizers. (Jonassen et al. 2021) provided the first proof of concept for this approach, demonstrating that organic residues from biogas plants were suitable as substrate and vector for cost-efficient, large-scale inoculation of NRBs to soil. While this proved effective in the short term, the organisms used were not suited to the soil environment and rapidly lost activity upon application. A second study by Jonassen et al. (2022) aimed at isolating NRB capable of competitive survival in both digestate and soil environments. To achieve this an enrichment procedure was designed involving sequential enrichment of a complex community, alternating between growth in sterilized soil and in digestate. Each enrichment was provided with ample amounts of N_2_O and a low initial O_2_ content (2 vol%), with O_2_ being depleted rapidly within the first 10-15 hours of the incubation. Conceptually, this *dual enrichment* selects for organisms able to grow both in digestate and soil, and with a capacity to adjust to rapid changes between the two substrates, as well as the rapid change between oxic and anoxic conditions. From these enrichments, number of N_2_O reducing organisms were isolated, one of which (*Cloacibacterium* sp. CB01) was particularly promising as it reached dominance in the enrichments and had a truncated denitrification pathway (NO and N_2_O reductases only).

16s rDNA amplicon sequencing showed that the dual substrate enrichment strategy effectively selected operational taxonomic units (OTUs) which were able to grow (or as a minimum survive) both in soil and digestate. The genetic elements possessed by and responsible for the success of these OTUs during the enrichment, however, remain unexamined. In the present work, we use combined metagenomics and metaproteomics on samples obtained from the enrichments carried out by Jonassen et al. (2022) to 1) examine the organisms and genes which were enriched during the dual substrate enrichment, with a focus on nitrogen transformation genes, 2) investigate what traits benefitted the enrichment winners, and 3) examine the proteins/metabolisms used by the enrichment winner (*Cloacibacterium* sp. CB01) during growth in the dual-substrate enrichment.

## Methods

### Enrichment culturing and sample collection

Samples for metagenomics and metaproteomics were taken from a dual-substrate enrichment culturing experiment described by Jonassen et al. (2022). Briefly, digestate was taken from an anaerobic digester at a municipal waste water treatment plant (Veas, Oslo, Norway) and clay loam soil from a long term liming experiment at the Norwegian University of Life Sciences (Nadeem et al. 2020). Enrichments were performed in 120 ml serum vials sealed with butyl rubber septa and incubated in a robotised incubation system at 20 °C (Molstad, Dörsch, and Bakken 2016; Molstad, Dörsch, and Bakken 2007). Headspace gas composition was repeatedly sampled and headspace gasses (O_2_, N_2_, N_2_O, NO, CO_2_ and CH_4_) quantified by gas chromatography (Molstad, Dörsch, and Bakken 2007). Initial enrichments were generated (n=7) using either 50 ml digestate (digestate enrichment series) or a mix of 20 g soil and 30 ml digestate (soil + digestate enrichment series). Headspace air was replaced with He then 3 ml N_2_O and 3 ml O_2_ added via injection. Vials were incubated at 20 °C with continuous stirring (600 rpm) for the digestate-containing vials. O_2_, N_2_O and N_2_ composition was monitored and additional N_2_O added to avoid depletion. Subsequent enrichments were then performed sequentially, switching between autoclaved digestate (121 °C for 20 mins then pH adjusted to 7.2 using HCl) and ϒ-irradiated soil (25.9 kGy, 12 months prior to use), as substrates. These enrichments were inoculated with 10 wt.% of the material from the preceding enrichment until 7 enrichment steps had been completed, each with 7 replicate vials.

### DNA isolation and sequencing

DNA was extracted from enrichment material in technical duplicates (1 ml digestate or 0.25 g soil) using the PowerLyzer™ Soil DNA extraction kit (QIAGEN) with modified bead beating using a MP Biomedicals™ FastPrep®-24 (Thermo Fischer Scientific Inc) for 30s at 4.5 ms-1 in place of vortexing. DNA from the seven replicate vials for each enrichment step were pooled for metagenomic sequencing. The TruSeq PCR-free library prep kit (illumina) was used to prepare a sequencing library followed by subsequent sequencing on an Illumina NovaSeq 6000 system with both 150 and 250 bp PE sequencing. Library preparation and sequencing were performed at The Norwegian Sequencing Centre (Oslo).

### Bioinformatics

Metagenomic sequence reads were trimmed with Trimmomatic V0.39. to remove adapter and low quality sequences (sliding window size 4 with minimum quality scores of 15 and minimum read length of 50bp), forward and reverse reads were merged where possible (Bolger, Lohse, and Usadel 2014).

#### Read based analyses

All forward, reverse and merged read files were concatenated for each sample for read based analyses. FMAP V0.15 was used to map reads to KEGG Orthology (KO) functional orthologs for read based functional analysis (Kim et al. 2016). Reads matching *nosZ* clade I and clade II genes were differentiated by aligning reads to a custom dataset of 21 *nosZ* clade I and 21 *nosZ* clade II protein sequences (Nadeau et al. 2019) using Diamond V2.0.9 where a maximum e-value was set to 1 x 10^-3^ and where all three top matches must match *nosZ* proteins of the same clade (Buchfink, Xie, and Huson 2015).

#### Assembly based metagenomic analyses

A co-assembly of trimmed reads from all samples was performed using Megahit V1.2.9 and contigs >1500 bp used for downstream analysis. (Li, Liu, et al. 2015). Reads from each sample were then mapped back against the assembled contigs using Bowtie2 V2.3.4.1 for the purpose of binning (Langmead and Salzberg 2012). Metagenomic binning of contigs was then performed using three binning software, Metabat2 V2.15 (Kang et al. 2019), Vamb V3.0.2 (Nissen et al. 2021), and maxbin2 V2.2.7 (Wu, Simmons, and Singer 2016). All three bin sets were then refined to a single set using Metawrap V1.2 (Uritskiy, DiRuggiero, and Taylor 2018) and CheckM V1.1.3 was used to assess the quality of these refined MAGs (Parks et al. 2015). Taxonomy of MAGs was assigned using GTDB-TK V1.5.1 (Chaumeil et al. 2020). MAGs were then compared to the genomes of organisms isolated from the enrichment, described in Jonassen et al (2022) (PRJEB44171) using OrthoANIu to match isolates with their corresponding MAG (Yoon et al. 2017). As no matches were found between the isolate genomes and any of the MAGs, isolate genomes were added to the MAG set and reads mapped against this dataset of MAGs and isolate genomes using Bowtie2 V2.3.4.1 before using CoverM V0.6.1 (https://github.com/wwood/CoverM) to assess the read coverage of the individual MAGs and isolates. This MAG set plus isolate genomes was then used for all subsequent analyses of MAGS. A phylogenetic tree was constructed for the HQ MAG set using GTOTREE v1.6.00 based on the pre-packaged set of 74 single copy bacterial genes (Lee 2019). Where at least 22 (30%) of the 74 single copy bacterial genes were required from a genome for inclusion. Briefly, this process included identification of genes using HMMER3 v3.2.2 (Eddy 2011), gene alignment with Muscle v5.1 (Edgar 2021), trimming with trimal v1.4.rev15 (Capella-Gutiérrez, Silla-Martínez, and Gabaldón 2009) and concatenation before phylogenetic tree building with FastTree2 v2.1.11 (Price, Dehal, and Arkin 2010). The resulting phylogenetic tree was then visualised using the Evolview3 webserver (Subramanian et al. 2019). Functional annotation was performed on the individual MAGs, isolates and the co-assembled contigs, separately, using PROKKA V1.14.6 (Seemann 2014). Kofamscan V1.3.0 (Aramaki et al. 2020) and KEGGdecoder V1.2.2 (Graham, Heidelberg, and Tully 2018) were then run on the PROKKA derived predicted protein coding sequences for each MAG to provide KEGG (Kyoto Encyclopaedia of Genes and Genomes) annotations. Gliding motility genes and Polysaccharide Utilisation Loci (PULs) were annotated for the two *Cloacibacterium* genomes (CB01 and CB03) together as distinguishing between the two was not possible for many of the proteins observed in the metaproteomics analysis. Gliding motility genes were annotated using BLASTp V2.12.0+ against a set of gliding motility proteins using an e-value cut-off of 0.00001 as recommended and presented by (Gavriilidou et al. 2020). PULs were identified and annotated using PULpy (Stewart et al. 2018).

### Quantitative metaproteomics

Samples from seven biological replicate enrichments were pooled and divided into three technical replicates for protein extraction. Thawed pooled samples were suspended in lysis buffer (20 mM Tris-HCl pH8, 0.1 % v/v Triton X-100, 200 mM NaCl, 1 mM DTT, 4% SDS, protease inhibitor cocktail (cOmplete, Mini, EDTA-free, Roche)) and treated with 3 × 45 s bead beating with glass beads (particle size ≤106 μm, Sigma) at maximum power and cooling on ice between cycles (MP Biomedicals™ FastPrep- 24TM, Thermo Fischer Scientific Inc). Cell debris and solid particles were removed by centrifugation (10 000 × *g*, 5 min). The protein concentration of lysates was determined by BCA protein assay (Thermo Fischer Scientific Inc) in order to normalize sample loadings prior to electrophoresis. The samples were loaded onto Any kD™ Mini-PROTEAN® TGX™ gels and separated under reducing conditions for 5 minutes at 250 V (Biorad). Proteins were visualized by Coomassie Blue Staining (Sigma). Each lane was cut into three fractions and corresponding fractions were pooled for each of the three replicates, reduced with 10 mM DTT for 30 min at 56°C, carbamidomethylated with 55 mM iodoacetamide for 30 min at room temperature, and further digested into peptides using 300 ng trypsin sample^-1^ at 37°C overnight. The peptides were desalted using C18 ZipTips (Merch Millipore, Darmstadt, Germany), according to manufacturer’s instructions.

The peptides were analysed by nanoLC-MS/MS using a Dionex Ultimate 3000 UHPLC (Thermo Fischer Scientific Inc) coupled to a Q-Exactive hybrid quadrupole orbitrap mass spectrometer (Thermo Scientific, Bremen, Germany). Peptides were separated using an analytical column (Acclaim PepMap RSLC C18, 2 µm, 100 Å, 75 µm i.d. × 50 cm, nanoViper) with a 120-minutes gradient from 3.2 to 44 % [v/v] acetonitrile in 0.1% [v/v] formic acid) at flow rate 300 nL/min. The Q-Exactive mass spectrometer was operated in data-dependent mode acquiring one full scan (400-1500 m/z) at R=70000 followed by (up to) 10 dependent MS/MS scans at R=35000.

The acquired MS/MS spectra were searched with MSFragger/FragPipe version 16 (Kong et al. 2017) against the metaproteome of either a focused (MAG-centric) database constructed from 290 MAGs (completeness >50%, contamination <20%) and recovered from the abovementioned metagenomics data (803 361 protein sequences), or a larger database generated from the same MAGs but also including unmapped contigs, i.e., contigs that could not be mapped to any of the MAGs (4 124 191 protein sequences). In addition, common contaminants such as human keratins, trypsin and bovine serum albumin were concatenated to the sample specific database as well as reversed sequences of all protein entries for estimation of false discovery rates. Proteins were quantified using the IonQuant algorithm in FragPipe (Yu, Haynes, and Nesvizhskii 2021). Protein N-terminal acetylation and oxidation of methionine were used as variable modifications, while carbamidomethylation of cysteine residues was used as a fixed modification. Tolerance levels for peptide identifications were ±20 ppm for both MS and MS/MS searches, and two missed cleavages of trypsin were allowed. All identifications were filtered to achieve a protein false discovery rate (FDR) of 1%. For a protein to be considered valid, we required the protein to be both identified and quantified in at least two of the three replicates.

## Results and discussion

Manipulation of the microbiomes of agricultural soils and the rhizosphere of crop plants is attractive because it could enhance the health and growth of crop plants (Oleńska et al. 2020), and degradation of pollutants (Saeed et al. 2021). In theory, this could be achieved by selective stimulation of indigenous microbes with desirable traits (Yuan et al 2021), but the most common approach has been to inoculate the systems with organisms with desirable traits such as plant-growth-promotion, suppression of pathogens, or degradation of pollutants. These attempts are often, however, unsuccessful in establishing a permanent or long-lasting presence of the introduced microbe in the target environment. The reasons for this are thought to stem from a combination of biotic and abiotic factors causing the inoculated microbe to fail to establish its own niche, whether it be metabolic or physical (Albright et al. 2022). Soil microbiomes in particular have been found to strongly repress any inoculants, thought to be due to their high species richness and diversity (van Elsas et al 2011). This failure of an inoculant to establish in a soil environment was observed by (Jonassen et al. 2021) when an NRB organism isolated from digestate was introduced to soil. Subsequently, an effort was made to obtain NRB’s with a better performance by including a selection pressure for growth, or at least survival, in soil. This was implemented by the dual substrate enrichment, whereby sequential N_2_O driven batch enrichments were performed, alternating between digestate and soil as substrates. In addition to selecting for the capacity to grow in both substrates, it plausibly also selected for organisms tolerant of rapid environmental changes: switching between soil and digestate several times, as well as an initial exposure to oxygen in each batch. The result of this was a set of isolates which could be grown in digestate but would maintain their activities, namely N_2_O reduction, when introduced to soil (Jonassen et al. 2022). Here, we investigate the genetic and proteomic basis for the success of this enrichment scheme by metagenomic and metaproteomic examination of the microbial community at each stage of the enrichment, and by analysing the genomes of the *Cloacibacterium* enrichment dominating isolates.

### Read based metagenomics and whole community metaproteomics – Enriched proteins

To investigate the genes and molecular functions which became more abundant throughout the enrichment culturing FMAP was used to map trimmed metagenomic reads to the KEGG Orthology database (supplementary material 1). This was then compared to the proteomics data which was sorted to identify the most increasingl proteins throughout the enrichment, thus examining which proteins that are increasingly being used by the microbial community as the enrichment progressed (Figure 1, supplementary material 1). It is noteworthy, as can be seen in Figure 1, that a large number of the genes that became abundant were not expressed. These genes were evidently carried by the organisms that reached dominance, but the proteins coded for were not in use. This highlights the importance of proteomic or transcriptomic approaches in interpreting the activity of microbial communities and confirms the observations that gene abundance often does not correlate with functional changes in microbial communities (Nannipieri et al. 2020; Nishisaka et al. 2019). Of the most increasingly expressed proteins throughout the enrichment three groups of proteins appear to be of particular importance here. Proteins involved in glycolysis, gluconeogenesis and the TCA cycle, environmental stress tolerance proteins, and motility, attachment and chemotaxis proteins (Figure 1).

**Figure 1.**
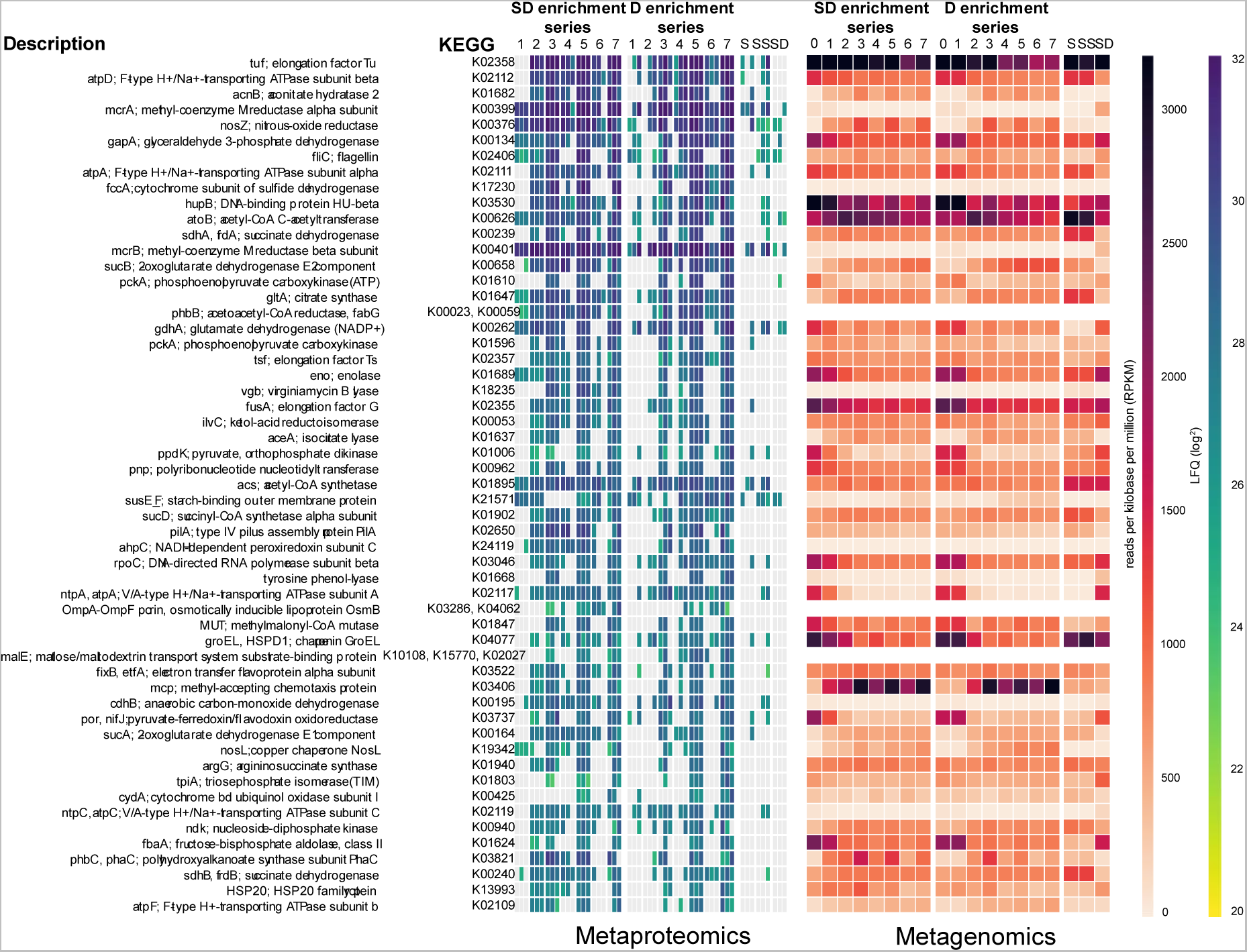
Metagenomic and metaproteomic quantification of the top 55 most increasing proteins in the dual substrate enrichments and their gene abundances. For each protein the slope of LFQ values from cycles 1 to 7 for both the D (digestate inoculated) and SD (soil + digestate inoculated) enrichment series was calculated and the average of the two slopes used to calculate the proteins with the greatest increases in abundance. Rows are ordered by slope from highest to lowest. Time point 0 for the metagenomic quantification refers to the read counts in the enrichment starting materials. For the metaproteomics heatmap each square box representing one enrichment is divided into three segments representing technical replicates. Samples S: soil, SS: gamma-sterilised soil, SD: autoclave sterilised digestate.

Of the top 55 most increasing proteins observed in the enrichment cultures, seven are involved in the glycolysis and gluconeogenesis pathways with one protein, fructose-bisphosphate aldolase (PpdK), specific to gluconeogenesis, indicating that this activity is not entirely a catabolic process but also anabolic. The presence of gluconeogenesis could be tied to biofilm formation (Tomlinson et al. 2021; Li, Chen, et al. 2015), which can aid an organism in establishing a physical niche and protecting it from environmental stressors such as changes in osmolarity or the presence of inhibitory compounds (Hall-Stoodley, Costerton, and Stoodley 2004). The production of glucose can also be used to produce glycogen or other storage carbohydrates which provide a buffer of carbon while the cells metabolism switches to utilisation of new carbon sources during rapid environmental change (Sekar et al. 2020). Also amongst the most increasing proteins in the enrichments were the proteins polyhydroxyalkanoate synthase subunit (PhbC) and acetoacetyl-CoA reductase (PhbB), responsible for the synthesis of polyhydroxyalkanoates, a carbon and energy storage molecule. Polyhydroxyalkanoates, like glycogen, have been demonstrated to provide increased survival rates during environmental stress conditions in a range of bacterial taxa (Castro-Sowinski et al. 2010).

Another interesting group of proteins observed to increase in abundance throughout the enrichment are those involved in motility, attachment and chemotaxis (Figure1). These include the flagellin protein (FliC) which is the major structural component from which a bacterial flagellum is composed, A methyl accepting chemotaxis protein (Mcp) and a type IV pilus assembly protein (PilA). Flagella provide motility, but can also play a major role in bacterial attachment to biotic (Haiko and Westerlund-Wikström 2013), and abiotic surfaces (Landini and Zehnder 2002), primarily by facilitating the initial attachment to surfaces (Conrad 2012). Similarly, type IV pili appear to facilitate initial attachment (Conrad 2012) as well as to secure motility along surfaces (Nudleman and Kaiser 2004; Pelicic 2008). This, taken with the observation of increasing quantities of methyl-accepting chemotaxis protein (Mcp) used for detecting chemical gradients, classically amino acids, and directing motility and attachment (Hida et al. 2020), suggests that the movement to physical niches and establishment there played an important role in survival and success in the dual substrate enrichment.

### Nitrogen cycle genes and proteins during enrichment

The number of metagenomic reads belonging to nitrogen cycling genes and the abundances of nitrogen cycling proteins was also traced throughout the enrichment process (Figure 2). As would be expected the number of *nosZ* reads increased throughout the enrichment, with a much greater enrichment of *nosZ* clade II genes than of clade I genes. This could be taken to suggest a selective pressure favouring organisms with NosZ clade II, but it cannot be excluded that the few nosZ clade II organisms that became dominant did so for other reasons than their nosZ clade II enzyme. *nirS* and both *norB* and *norC* appeared to be co-enriched while minimal enrichment of nitrate reductase genes (Nap, Nar) was seen, indicating the enrichment of truncated denitrifiers over those with a fully-fledged denitrification system. Regarding the expression of nitrogen cycling enzyme expression, nosZ was abundant throughout, NarG, NarH and NorC were observed in the initial enrichment but were subsequently lost, while NirS and to a lesser extent NirK were persistently present throughout the enrichments, despite the fact that no nitrate or nitrite was added. The gamma-sterilized soil contained traces of NO_2_ ^-^, however and during the first 15 hours of enrichment culturing in soil, NO accumulated transiently to 4-6 nM in the liquid. In contrast, the digestate enrichment cultures showed a marginal but constant rate of NO accumulation throughout, reaching 0.5-0.7 nM at the end of each incubation (supplementary material 2).

**Figure 2.**
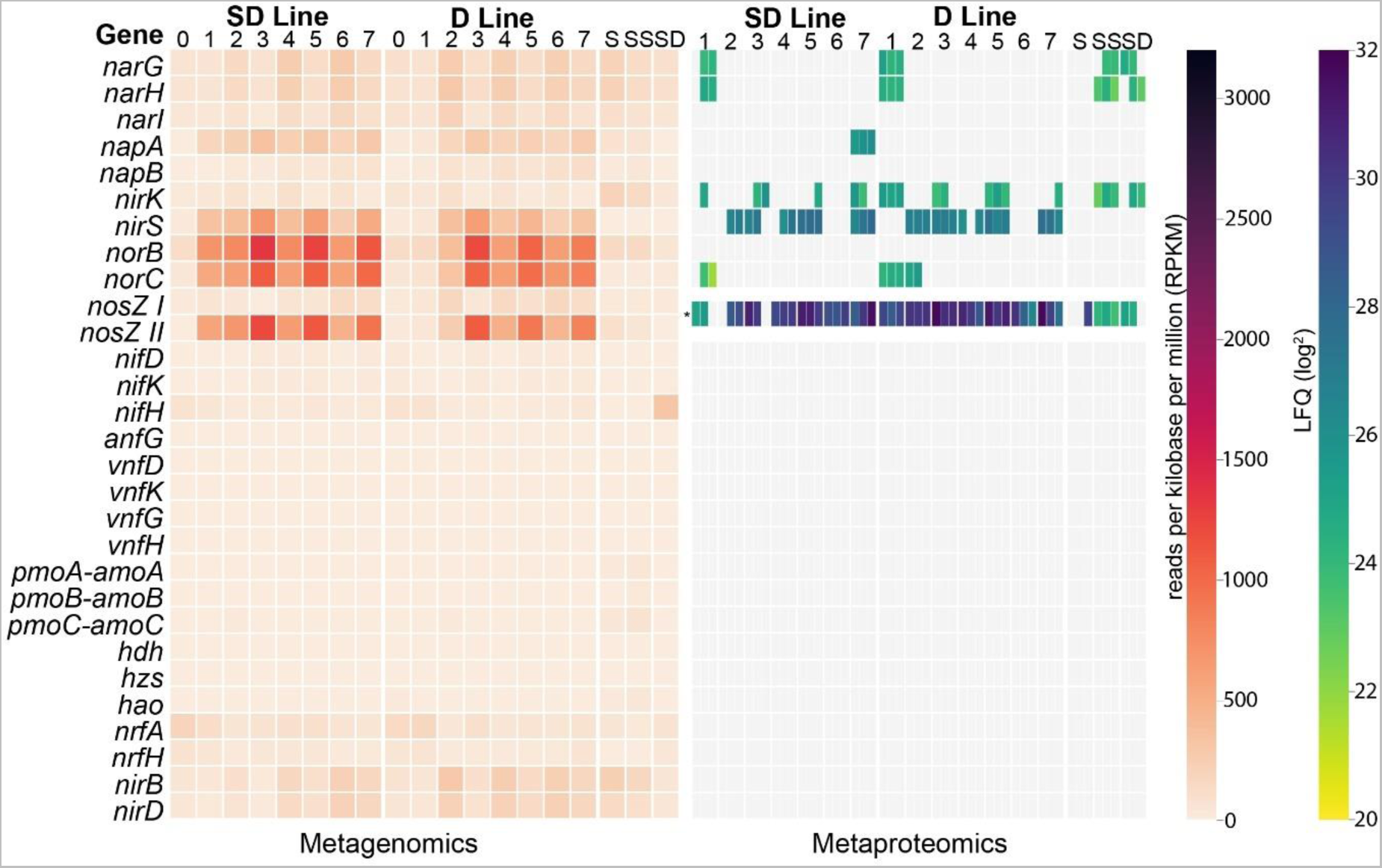
Metagenomic and metaproteomic quantification of nitrogen cycling genes and proteins in the dual substrate enrichment. Enrichment step zero for the metagenomic quantification refers to the first enrichment culture, immediately after inoculation. *There was insufficient sequence information to distinguish between NosZ clade I and II proteins, so they are combined for the proteomic analysis. Samples S: soil, SS: gamma-sterilised soil, SD: autoclave sterilised digestate.

### Genome-centric metagenomics

While read based analysis of metagenomics datasets can give good quantitative information on the functional genes and inform on their enrichment or loss from various samples, the taxonomic affiliations and genomic contexts of these genes, and their corresponding proteins, are difficult to infer. For this reason, a genome-centric analysis of the metagenomic data was undertaken to group metagenomic reads into the genomes from which they originate. Genome binning of the metagenomic reads from the current study resulted in the generation of 290 high-quality MAGs. MAGs were compared against the isolate genomes obtained from the enrichment experiment (Jonassen et al. 2022) and where there was no match to any MAG those genomes were added to the MAG set, as metagenomic binning had failed to identify these genomes. Amongst these high-quality MAGs and isolate genomes, 22 were found to contain the *nosZ* gene, with the majority seen in the Proteobacteria. Amongst these MAGs and genomes there were very few which appeared to increase in abundance throughout the enrichment with the notable exception of the two *Cloacibacterium* isolate genomes. These matched the single OTU identified by Jonassen et al. (2022) as the enrichment winning organism. The relative abundances of these two organisms, however, were lower than the abundance of the OTU identified by Jonassen et al. (2022), suggesting that additional closely related *Cloacibacterium* organisms are likely to be present, circumscribed by the 16S OTU, but not mapping to the isolate genomes or assembling into MAGs (Figure 3). Additionally, the 16S gene copy number is three in the *Cloacibacterium* genomes which may differ from the average for the community skewing the observed abundance in the 16S rDNA amplicon data.

**Figure 3:**
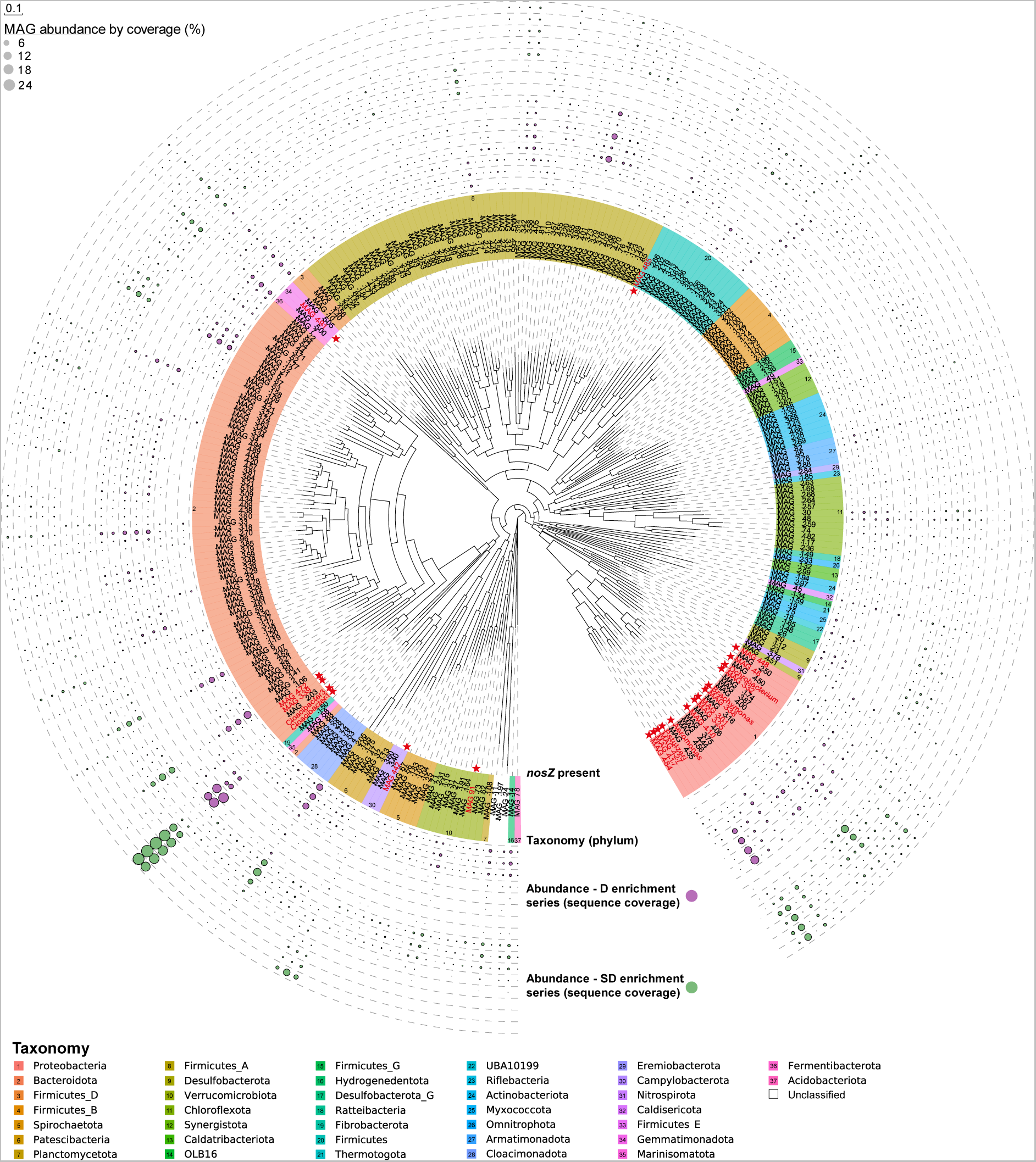
MAGS and isolate genomes from dual enrichment culturing metagenomes. A phylogenetic maximum likelihood tree of the high-quality MAGs and isolates’ genomes retrieved from the dual enrichment culturing metagenomes based a set of concatenated single-copy gene translated protein sequences. Taxonomy at the phylum level was assigned using the GTDBTK and is indicated by leaf node colour. The relative abundances of the MAGs and isolate genomes as calculated by MAG metagenomic read coverage in the various samples are indicated by bubbles at the branch tips for both the SD (soil + digestate inoculated) (green) and D (digestate inoculated) (purple) enrichment series. The detection of a *nosZ* gene in the MAG or isolate genome is indicated by red colouring of the MAG label and a red star icon. Thirteen of the 294 high-quality MAGs did not include enough single-copy gene sequences for inclusion in the phylogenetic tree and are not shown.

### Genome-centric metaproteomics of *Cloacibacterium*

In order to examine the metabolism and activities of the enrichment winning *Cloacibacterium* organisms, a genome-centric analysis of the metaproteomics was performed by creating a proteomics reference database from the MAGs plus the isolates’ genomes. This allowed for the proteins observed in the metaproteomics to be ascribed to individual MAGs or genomes, and their abundances traced throughout enrichment cycles. Due to the similarities in the genes held by the two *Cloacibacterium* isolate genomes many of the proteins identified by metaproteomics could not be distinguished between the two organisms and so the metaproteomic analysis of these two organisms is combined (Figure 4). A large number of Carbohydrate Active Enzymes (CAZymes) was identified in the *Cloacibacterium* proteomes throughout the enrichment, with several identified to be part of Polysaccharide Utilisation Loci (PULs). These PULs are specific to the Bacteroidetes and are modular colocalized gene clusters encoding recognition, binding, transport and enzymatic proteins. Each cluster typically contains a gene suit necessary for the targeted recognition, breakdown and import of a specific carbohydrate. Recognition and carbohydrate cleavage enzymes, termed glycoside hydrolases (GHs), are located at the cell surface where they bind and carry out the initial stages of carbohydrate degradation, followed by the binding and import of the smaller carbohydrate products into the cell’s periplasm and then cytoplasm. Each PUL can contain several glycoside hydrolases and the type and combination of glycoside hydrolases determines the substrate specificity (Grondin et al. 2017). PULs are normally tightly regulated, only expressed when their substrates are detected in abundance (Larsbrink and McKee 2020). The *Cloacibacterium* genomes contained nine distinct PULs, seven of which contained proteins which were observed by metaproteomics and in some cases could be closely enough matched to characterised PULs in the Polysaccharide-Utilization Loci DataBase (PULDB) to identify their likely substrates (Terrapon et al. 2018). PUL2 contains genes from the glycoside hydrolase enzyme families GH13 and GH97 which can hydrolyse a wide range of α-glucoside and α-galactoside linkages and so have a broad range of substrates (Gloster et al. 2008; Mewis et al. 2016). PUL4 and PUL8 possess GH30 and GH3 family enzymes which also have various activities, including β-1,6-glucanase, and β-xylosidase activities. PULs with these two GH families have been reported to be associated with the degradation of plant and fungal beta-1,6-glucans or xylose (Martens et al. 2011; Temple et al. 2017; McBride et al. 2009). PUL6 contains the GH2, GH13, GH53 and GH97 family enzymes, many of which target multiple substrates. GH53 is relatively specific and only has known activity associated with the breakdown galactans and arabinogalactans in the pectic component of plant cell walls (Luis et al. 2018; Hinz et al. 2005; Braithwaite et al. 1997) (Figure 4). The presence of these PULs and their expression during enrichment indicates the importance of complex carbohydrate degradation by *Cloacibacterium* during enrichment, which plausibly reflects that complex carbohydrates are one of the remaining available carbon sources not entirely depleted during the foregoing anaerobic digestion (biogas production). To complement the suit of PUL associated genes, a high number of glycolysis proteins were observed from *Cloacibacterium* in the enrichment, strengthening support for carbohydrate driven metabolism in these organisms during the enrichment culturing (Figure 4). Enzymes for synthesis and degradation of glycogen were also observed, suggesting that this system for surviving feast/famine periods was important for *Cloacibacterium* during the enrichment culturing, as was also suggested by the read based analysis above (Figures 1 and 4) (Sekar et al. 2020).

**Figure 4.**
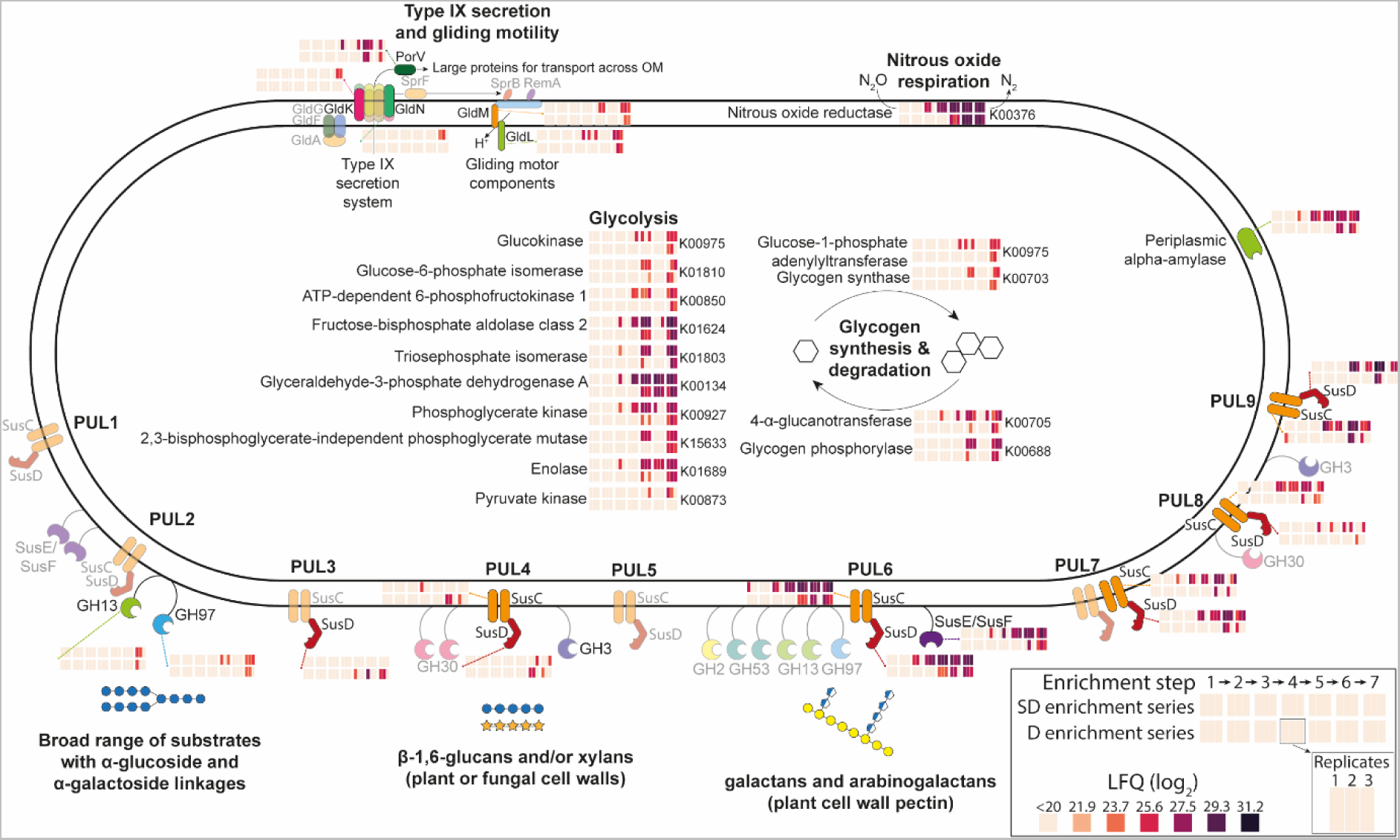
Diagram of genes and proteins expressed by *Cloacibacterium* CB01 and CB03 during enrichment culturing. For each protein, the abundance in the enrichments 1-7 (from left to right) are shown as heat maps, one box for each enrichment, divided in three segments representing technical replicates. The upper row shows the result for the SD enrichment series (inoculated with soil and digestate), the lower row for the D enrichment series (inoculated with digestate only). Opaque shapes and protein names denote proteins for which a coding gene was present in the genome but not observed in the metaproteomics.

Another component of the *Cloacibacterium* genomes for which there was proteomic evidence during enrichment was the type IX secretion system (T9SS) and gliding motility machinery (Figure 4). The T9SS is a complex multicomponent secretion system unique to the Fibrobacteres-Chlorobi-Bacteroidetes superphylum, used primarily for the export of proteins destined to be anchored to the cell surface. Proteins excreted by the type IX secretion system include enzymes involved in the breakdown of complex polymers, including those encoded by the PUL systems, and the gliding motility machinery which is also unique to this taxonomic group (McKee et al. 2021). Two proteins from the gliding motility apparatus were observed from the *Cloacibacterium* in the enrichment culturing, GldM and GldL (Figure 4). These proteins make up the stator and rotor in a membrane bound molecular motor which drives the movement of adhesins across the cell surface, facilitating cell gliding across a solid substrate (Veith et al. 2017; McBride 2019). The expression of the gliding motility system would allow *Cloacibacterium* to migrate and attach to insoluble polysaccharides within the enrichment medium whereby the relevant PULs would be expressed, and their degradative enzymes transported to the cell surface to degrade the substrate. Gliding motility genes have also been shown to be involved in biofilm formation in another member of the Bacteroides (Sato et al. 2021). This trait would aid in maintaining localisation within a desirable micro-niche.

## Conclusions

The discovery of isolates capable of modifying a native microbial community in order to introduce new traits has long been of interest to diverse biotechnologists but has often come up against issues of poor invasion success (Van Elsas et al. 2012; Han et al. 2021). The recent work of Jonassen et al. (2022) has focussed on this issue in their attempt to isolate N_2_O reducing organisms for introduction to soils via organic wastes as a substrate and vector, by utilising a *dual enrichment* procedure designed to enrich organisms with a high tolerance for a rapidly changing environment. In the current study the genetic and proteomic factors behind the success of this enrichment have been explored. Through a read-based and genome-centric analysis the factors and strategies which appear to be utilised by enrichment winners during the enrichment were identified and include complex carbohydrate metabolism, motility and attachment (biofilm, gliding motility, chemotaxis, type IV pili) and the production of energy and carbon storage molecules used in a feast/famine lifestyle (glycogen and polyhydroxyalkanoates). This was true for the major *Cloacibacterium* enrichment winners as well as other organisms not identified by metagenomic genome binning (Figures 1 and 4). From a community perspective *nosZ* was co-enriched with nitrite reductase and nitric oxide reductase genes but not nitrate reductase genes. Additionally, the two isolated enrichment winners were truncated denitrifiers; one a norBC + *nosZ* organism and the other a norBC + *nosZ* + *napA* carrying organism. While *Nor* and *Nir* (S and K) genes appeared to be enriched, only NirS was expressed as seen in the proteomics throughout enrichment.

An omics-based perspective of this dual enrichment culturing procedure has provided a better understanding of the mechanisms used by denitrifying organisms to tolerate and proliferate under rapid environmental changes as well as a better understanding of the enrichment winning *Cloacibacterium* isolates and what makes them so successful as inoculants to soils where they are able to persist and maintain N_2_O reductase activity (Jonassen et al. 2022). These findings could be used to optimise the application of *Cloacibacterium* as a soil inoculant for modifying the N_2_O reduction potential of soils, including the use of altered carrier materials rich in plant based complex polysaccharides which can also provide a physical micro-niche from soil competitors and nosZ inhibiting low pH. These findings are also of interest for researchers interested in the barriers to establishment of microbes in new environments with an existing microbial community.

## Data Availability

Metagenomic reads have been uploaded to the Sequence read Archive (SRA) and are available within the BioProject PRJNA909336. The mass spectrometry proteomics data have been deposited to the ProteomeXchange Consortium via the PRIDE (Perez-Riverol et al. 2022) partner repository with the dataset identifier PXD037372.

## Acknowledgements

Mass spectrometry-based proteomic analyses were performed by the MS and Proteomics Core Facility, Norwegian University of Life Sciences (NMBU). This facility is a member of the National Network of Advanced Proteomics Infrastructure (NAPI), which is funded by the Research Council of Norway INFRASTRUKTUR-program (project number: 295910). The authors acknowledge the Orion High Performance Computing Center (OHPCC) at the Norwegian University of Life Sciences (NMBU) for providing computational resources that have contributed to the research results reported within this paper. This work was financially supported by the Norwegian Research Council (project number 260868).

## Notes

### Competing Interest Statement

The authors have declared no competing interest.

## References

Albright, Michaeline BN, Stilianos Louca, Daniel E Winkler, Kelli L Feeser, Sarah-Jane Haig, Katrine L Whiteson, Joanne B Emerson, and John Dunbar. 2022. ’Solutions in microbiome engineering: prioritizing barriers to organism establishment’, The ISME Journal, 16: 331–38.

Aramaki, Takuya, Romain Blanc-Mathieu, Hisashi Endo, Koichi Ohkubo, Minoru Kanehisa, Susumu Goto, and Hiroyuki Ogata. 2020. ’KofamKOALA: KEGG ortholog assignment based on profile HMM and adaptive score threshold’, Bioinformatics, 36: 2251–52.

Bolger, Anthony M, Marc Lohse, and Bjoern Usadel. 2014. ’Trimmomatic: a flexible trimmer for Illumina sequence data’, Bioinformatics, 30: 2114–20.

Braithwaite, Kerynne L, Tereza Barna, Tracey D Spurway, Simon J Charnock, Gary W Black, Neil Hughes, Jeremy H Lakey, Richard Virden, Geoffrey P Hazlewood, and Bernard Henrissat. 1997. ’Evidence that galactanase A from Pseudomonas fluorescens subspecies cellulosa is a retaining family 53 glycosyl hydrolase in which E161 and E270 are the catalytic residues’, Biochemistry, 36: 15489–500.

Buchfink, Benjamin, Chao Xie, and Daniel H Huson. 2015. ’Fast and sensitive protein alignment using DIAMOND’, Nature methods, 12: 59–60.

Butterbach-Bahl, Klaus, Elizabeth M Baggs, Michael Dannenmann, Ralf Kiese, and Sophie Zechmeister-Boltenstern. 2013a. ’Nitrous oxide emissions from soils: how well do we understand the processes and their controls?’, Philosophical Transactions of the Royal Society B: Biological Sciences, 368: 20130122.

Butterbach-Bahl, Klaus, Elizabeth M Baggs, Michael Dannenmann, Ralf Kiese, and Sophie Zechmeister-Boltenstern. 2013b. ’Nitrous oxide emissions from soils: how well do we understand the processes and their controls?’, Philosophical Transactions of the Royal Society B: Biological Sciences, 368.

Capella-Gutiérrez, Salvador, José M Silla-Martínez, and Toni Gabaldón. 2009. ’trimAl: a tool for automated alignment trimming in large-scale phylogenetic analyses’, Bioinformatics, 25: 1972–73.

Castro-Sowinski, Susana, Saul Burdman, Ofra Matan, and Yaacov Okon. 2010. ’Natural functions of bacterial polyhydroxyalkanoates.’ in, Plastics from bacteria (Springer).

Chaumeil, Pierre-Alain, Aaron J Mussig, Philip Hugenholtz, and Donovan H Parks. 2020. ’GTDB-Tk: a toolkit to classify genomes with the Genome Taxonomy Database’, Bioinformatics, 36: 1925– 27.

Conrad, Jacinta C. 2012. ’Physics of bacterial near-surface motility using flagella and type IV pili: implications for biofilm formation’, Research in Microbiology, 163: 619–29.

Davidson, Eric A. 2009. ’The contribution of manure and fertilizer nitrogen to atmospheric nitrous oxide since 1860’, Nature Geoscience, 2: 659–62.

Domeignoz-Horta, Luiz A, Martina Putz, Aymé Spor, David Bru, Marie-Christine Breuil, Sara Hallin, and Laurent Philippot. 2016. ’Non-denitrifying nitrous oxide-reducing bacteria-An effective N2O sink in soil’, Soil Biology and Biochemistry, 103: 376–79.

Eddy, Sean R. 2011. ’Accelerated profile HMM searches’, PLoS Computational Biology, 7: e1002195.

Edgar, Robert C. 2021. ’MUSCLE v5 enables improved estimates of phylogenetic tree confidence by ensemble bootstrapping’, bioRxiv.

Gao, Nan, Weishou Shen, Estefania Camargo, Yutaka Shiratori, Tomoyasu Nishizawa, Kazuo Isobe, Xinhua He, and Keishi Senoo. 2017. ’Nitrous oxide (N 2 O)-reducing denitrifier-inoculated organic fertilizer mitigates N 2 O emissions from agricultural soils’, Biology and Fertility of Soils, 53: 885–98.

Gavriilidou, Asimenia, Johanna Gutleben, Dennis Versluis, Francesca Forgiarini, Mark WJ van Passel, Colin J Ingham, Hauke Smidt, and Detmer Sipkema. 2020. ’Comparative genomic analysis of Flavobacteriaceae: Insights into carbohydrate metabolism, gliding motility and secondary metabolite biosynthesis’, Bmc Genomics, 21: 1–21.

Gloster, Tracey M, Johan P Turkenburg, Jennifer R Potts, Bernard Henrissat, and Gideon J Davies. 2008. ’Divergence of catalytic mechanism within a glycosidase family provides insight into evolution of carbohydrate metabolism by human gut flora’, Chemistry and Biology, 15: 1058–67.

Graham, ED, JF Heidelberg, and BJ Tully. 2018. ’Potential for primary productivity in a globally-distributed bacterial phototroph’, The ISME journal, 12: 1861–66.

Grondin, Julie M, Kazune Tamura, Guillaume Déjean, D Wade Abbott, and Harry Brumer. 2017. ’Polysaccharide utilization loci: fueling microbial communities’, Journal of Bacteriology, 199: e00860–16.

Haiko, Johanna, and Benita Westerlund-Wikström. 2013. ’The role of the bacterial flagellum in adhesion and virulence’, Biology, 2: 1242–67.

Hall-Stoodley, Luanne, J William Costerton, and Paul Stoodley. 2004. ’Bacterial biofilms: from the natural environment to infectious diseases’, Nature Reviews Microbiology, 2: 95–108.

Hallin, Sara, Laurent Philippot, Frank E Löffler, Robert A Sanford, and Christopher M Jones. 2018. ’Genomics and ecology of novel N2O-reducing microorganisms’, Trends in Microbiology, 26: 43–55.

Han, Shengyi, Yanmeng Lu, Jiaojiao Xie, Yiqiu Fei, Guiwen Zheng, Ziyuan Wang, Jie Liu, Longxian Lv, Zongxin Ling, and Björn Berglund. 2021. ’Probiotic gastrointestinal transit and colonization after oral administration: A long journey’, Frontiers in Cellular and Infection Microbiology, 11: 609722.

Hida, Akiko, Shota Oku, Manami Miura, Hiroki Matsuda, Takahisa Tajima, and Junichi Kato. 2020. ’Characterization of methyl-accepting chemotaxis proteins (MCPs) for amino acids in plant-growth-promoting rhizobacterium Pseudomonas protegens CHA0 and enhancement of amino acid chemotaxis by MCP genes overexpression’, Bioscience, Biotechnology, and Biochemistry, 84: 1948–57.

Hinz, Sandra WA, Marieke I Pastink, Lambertus AM van den Broek, Jean-Paul Vincken, and Alphons GJ Voragen. 2005. ’Bifidobacterium longum endogalactanase liberates galactotriose from type I galactans’, Applied and Environmental Microbiology, 71: 5501–10.

Jonassen, Kjell Rune, Live H Hagen, Silas HW Vick, Magnus Ø Arntzen, Vincent GH Eijsink, Åsa Frostegård, Pawel Lycus, Lars Molstad, Phillip B Pope, and Lars R Bakken. 2021. ’Nitrous oxide respiring bacteria in biogas digestates for reduced agricultural emissions’, The ISME Journal: 1–11.

Jonassen, Kjell Rune, Ida Ormåsen, Clara Duffner, Torgeir R Hvidsten, Lars R Bakken, and Silas HW Vick. 2022. ’A Dual Enrichment Strategy Provides Soil-and Digestate-Competent Nitrous Oxide-Respiring Bacteria for Mitigating Climate Forcing in Agriculture’, mBio: e00788–22.

Kang, Dongwan D, Feng Li, Edward Kirton, Ashleigh Thomas, Rob Egan, Hong An, and Zhong Wang. 2019. ’MetaBAT 2: an adaptive binning algorithm for robust and efficient genome reconstruction from metagenome assemblies’, PeerJ, 7: e7359.

Kim, Jiwoong, Min Soo Kim, Andrew Y Koh, Yang Xie, and Xiaowei Zhan. 2016. ’FMAP: functional mapping and analysis pipeline for metagenomics and metatranscriptomics studies’, BMC bioinformatics, 17: 1–8.

Kong, Andy T, Felipe V Leprevost, Dmitry M Avtonomov, Dattatreya Mellacheruvu, and Alexey I Nesvizhskii. 2017. ’MSFragger: ultrafast and comprehensive peptide identification in mass spectrometry–based proteomics’, Nature methods, 14: 513–20.

Landini, Paolo, and Alexander JB Zehnder. 2002. ’The global regulatory hns gene negatively affects adhesion to solid surfaces by anaerobically grown Escherichia coli by modulating expression of flagellar genes and lipopolysaccharide production’, Journal of Bacteriology, 184: 1522–29.

Langmead, Ben, and Steven L Salzberg. 2012. ’Fast gapped-read alignment with Bowtie 2’, Nature methods, 9: 357–59.

Larsbrink, Johan, and Lauren Sara McKee. 2020. ’Bacteroidetes bacteria in the soil: Glycan acquisition, enzyme secretion, and gliding motility’, Advances in Applied Microbiology, 110: 63–98.

Lee, Michael D. 2019. ’GToTree: a user-friendly workflow for phylogenomics’, Bioinformatics, 35: 4162–64.

Li, Dinghua, Chi-Man Liu, Ruibang Luo, Kunihiko Sadakane, and Tak-Wah Lam. 2015. ’MEGAHIT: an ultra-fast single-node solution for large and complex metagenomics assembly via succinct de Bruijn graph’, Bioinformatics, 31: 1674–76.

Li, Zhenjian, Yong Chen, Dong Liu, Nan Zhao, Hao Cheng, Hengfei Ren, Ting Guo, Huanqing Niu, Wei Zhuang, and Jinglan Wu. 2015. ’Involvement of glycolysis/gluconeogenesis and signaling regulatory pathways in Saccharomyces cerevisiae biofilms during fermentation’, Frontiers in microbiology, 6: 139.

Luis, Ana S, Jonathon Briggs, Xiaoyang Zhang, Benjamin Farnell, Didier Ndeh, Aurore Labourel, Arnaud Baslé, Alan Cartmell, Nicolas Terrapon, and Katherine Stott. 2018. ’Dietary pectic glycans are degraded by coordinated enzyme pathways in human colonic Bacteroides’, Nature Microbiology, 3: 210–19.

Lycus, Pawel, Kari Lovise Bøthun, Linda Bergaust, James Peele Shapleigh, Lars Reier Bakken, and Åsa Frostegård. 2017. ’Phenotypic and genotypic richness of denitrifiers revealed by a novel isolation strategy’, The ISME Journal, 11: 2219–32.

Martens, Eric C, Elisabeth C Lowe, Herbert Chiang, Nicholas A Pudlo, Meng Wu, Nathan P McNulty, D Wade Abbott, Bernard Henrissat, Harry J Gilbert, and David N Bolam. 2011. ’Recognition and degradation of plant cell wall polysaccharides by two human gut symbionts’, PLoS Biology, 9: e1001221.

McBride, Mark J. 2019. ’Bacteroidetes gliding motility and the type IX secretion system’, Microbiology Spectrum, 7: 7.1. 15.

McBride, Mark J, Gary Xie, Eric C Martens, Alla Lapidus, Bernard Henrissat, Ryan G Rhodes, Eugene Goltsman, Wei Wang, Jian Xu, and David W Hunnicutt. 2009. ’Novel features of the polysaccharide-digesting gliding bacterium Flavobacterium johnsoniae as revealed by genome sequence analysis’, Applied and Environmental Microbiology, 75: 6864–75.

McKee, Lauren S, Sabina Leanti La Rosa, Bjørge Westereng, Vincent G Eijsink, Phillip B Pope, and Johan Larsbrink. 2021. ’Polysaccharide degradation by the Bacteroidetes: mechanisms and nomenclature’, Environmental Microbiology Reports, 13: 559–81.

Mewis, Keith, Nicolas Lenfant, Vincent Lombard, and Bernard Henrissat. 2016. ’Dividing the large glycoside hydrolase family 43 into subfamilies: a motivation for detailed enzyme characterization’, Applied and Environmental Microbiology, 82: 1686–92.

Molstad, Lars, Peter Dörsch, and L Bakken. 2016. “Improved robotized incubation system for gas kinetics in batch cultures.” In.: Technical report.

Molstad, Lars, Peter Dörsch, and Lars R Bakken. 2007. ’Robotized incubation system for monitoring gases (O2, NO, N2O N2) in denitrifying cultures’, Journal of Microbiological Methods, 71: 202–11.

Nadeau, Sarah A, Constance A Roco, Spencer J Debenport, Todd R Anderson, Kathryn L Hofmeister, M Todd Walter, and James P Shapleigh. 2019. ’Metagenomic analysis reveals distinct patterns of denitrification gene abundance across soil moisture, nitrate gradients’, Environmental microbiology, 21: 1255–66.

Nadeem, Shahid, Lars R Bakken, Åsa Frostegård, John C Gaby, and Peter Dörsch. 2020. ’Contingent Effects of Liming on N2O-Emissions Driven by Autotrophic Nitrification’, Frontiers in Environmental Science, 8.

Nannipieri, Paolo, Judith Ascher-Jenull, Maria Teresa Ceccherini, Giacomo Pietramellara, Giancarlo Renella, and Michael Schloter. 2020. ’Beyond microbial diversity for predicting soil functions: A mini review’, Pedosphere, 30: 5–17.

Nishisaka, Caroline Sayuri, Connor Youngerman, Laura K Meredith, Janaina Braga do Carmo, and Acacio Aparecido Navarrete. 2019. ’Differences in N2O fluxes and denitrification gene abundance in the wet and dry seasons through soil and plant residue characteristics of tropical tree crops’, Frontiers in Environmental Science, 7: 11.

Nissen, Jakob Nybo, Joachim Johansen, Rosa Lundbye Allesøe, Casper Kaae Sønderby, Jose Juan Almagro Armenteros, Christopher Heje Grønbech, Lars Juhl Jensen, Henrik Bjørn Nielsen, Thomas Nordahl Petersen, and Ole Winther. 2021. ’Improved metagenome binning and assembly using deep variational autoencoders’, Nature biotechnology, 39: 555–60.

Nudleman, Eric, and Dale Kaiser. 2004. ’Pulling together with type IV pili’, Microbial Physiology, 7: 52–62.

Oleńska, Ewa, Wanda Małek, Małgorzata Wójcik, Izabela Swiecicka, Sofie Thijs, and Jaco Vangronsveld. 2020. ’Beneficial features of plant growth-promoting rhizobacteria for improving plant growth and health in challenging conditions: A methodical review’, Science of the Total Environment, 743: 140682.

Parks, Donovan H, Michael Imelfort, Connor T Skennerton, Philip Hugenholtz, and Gene W Tyson. 2015. ’CheckM: assessing the quality of microbial genomes recovered from isolates, single cells, and metagenomes’, Genome research, 25: 1043–55.

Pelicic, Vladimir. 2008. ’Type IV pili: e pluribus unum?’, Molecular Microbiology, 68: 827–37.

Perez-Riverol, Yasset, Jingwen Bai, Chakradhar Bandla, David García-Seisdedos, Suresh Hewapathirana, Selvakumar Kamatchinathan, Deepti J Kundu, Ananth Prakash, Anika Frericks-Zipper, and Martin Eisenacher. 2022. ’The PRIDE database resources in 2022: a hub for mass spectrometry-based proteomics evidences’, Nucleic Acids Research, 50: D543–D52.

Philippot, Laurent, Janet Andert, Christopher M Jones, David Bru, and Sara Hallin. 2011. ’Importance of denitrifiers lacking the genes encoding the nitrous oxide reductase for N2O emissions from soil’, Global Change Biology, 17: 1497–504.

Price, Morgan N, Paramvir S Dehal, and Adam P Arkin. 2010. ’FastTree 2–approximately maximum-likelihood trees for large alignments’, PloS One, 5: e9490.

Robertson, GP. 2014. ’Soil greenhouse gas emissions and their mitigation.’ in N. Van Alfen (ed.), Encyclopedia of agriculture and food systems (Elsevier: San Diego, California, USA).

Saeed, Qudsia, Wang Xiukang, Fasih Ullah Haider, Jiří Kučerik, Muhammad Zahid Mumtaz, Jiri Holatko, Munaza Naseem, Antonin Kintl, Mukkaram Ejaz, and Muhammad Naveed. 2021. ’Rhizosphere bacteria in plant growth promotion, biocontrol, and bioremediation of contaminated sites: a comprehensive review of effects and mechanisms’, International journal of molecular sciences, 22: 10529.

Sato, Keiko, Masami Naya, Yuri Hatano, Yoshio Kondo, Mari Sato, Keiji Nagano, Shicheng Chen, Mariko Naito, and Chikara Sato. 2021. ’Biofilm spreading by the adhesin-dependent gliding motility of Flavobacterium johnsoniae. 1. Internal structure of the biofilm’, International journal of molecular sciences, 22: 1894.

Seemann, Torsten. 2014. ’Prokka: rapid prokaryotic genome annotation’, Bioinformatics, 30: 2068–69.

Sekar, Karthik, Stephanie M Linker, Jen Nguyen, Alix Grünhagen, Roman Stocker, and Uwe Sauer. 2020. ’Bacterial glycogen provides short-term benefits in changing environments’, Applied and Environmental Microbiology, 86: e00049–20.

Shapleigh, James P. 2013. ’Denitrifying Prokaryotes.’ in Eugene Rosenberg, Edward F. DeLong, Stephen Lory, Erko Stackebrandt and Fabiano Thompson (eds.), The Prokaryotes: Prokaryotic Physiology and Biochemistry (Springer Berlin Heidelberg: Berlin, Heidelberg).

Solomon, S, Qin D, Manning M, Alley R.B, Berntsen T, Bindoff N.L, Chen Z, Chidthaisong A, Gregory J.M, Hegerl G.C, Heimann M, Hewitson B, Hoskins B.J, Joos F, Jouzel J, Kattsov V, Lohmann U, Matsuno T, Molina M, Nicholls N, Overpeck J, Raga G, Ramaswamy V, Ren J, Rusticucci M, Somerville R, Stocker T.F, Whetton P, Wood R.A, and Wratt D. 2007. ’Technical Summary.’ in

S. Solomon, D. Qin, M. Manning, Z. Chen, M. Marquis, K.B. Averyt, M. Tignor and H.L. Miller (eds.), Climate Change 2007: The Physical Science Basis. Contribution of Working Group I to the Fourth Assessment Report of the Intergovernmental Panel on Climate Change (Cambridge University Press: Cambridge, United Kingdom and New York, NY USA).

Stewart, Rob D., Marc D. Auffret, Rainer Roehe, and Mick Watson. 2018. ’Open prediction of polysaccharide utilisation loci (PUL) in 5414 public <em>Bacteroidetes</em> genomes using PULpy’, bioRxiv: 421024.

Subramanian, Balakrishnan, Shenghan Gao, Martin J Lercher, Songnian Hu, and Wei-Hua Chen. 2019. ’Evolview v3: a webserver for visualization, annotation, and management of phylogenetic trees’, Nucleic acids research, 47: W270–W75.

Temple, Max J, Fiona Cuskin, Arnaud Baslé, Niall Hickey, Gaetano Speciale, Spencer J Williams, Harry J Gilbert, and Elisabeth C Lowe. 2017. ’A Bacteroidetes locus dedicated to fungal 1, 6-β- glucan degradation: unique substrate conformation drives specificity of the key endo-1, 6-β- glucanase’, Journal of Biological Chemistry, 292: 10639–50.

Terrapon, Nicolas, Vincent Lombard, Elodie Drula, Pascal Lapébie, Saad Al-Masaudi, Harry J Gilbert, and Bernard Henrissat. 2018. ’PULDB: the expanded database of Polysaccharide Utilization Loci’, Nucleic Acids Research, 46: D677–D83.

Thompson, RL, L Lassaletta, PK Patra, C Wilson, KC Wells, A Gressent, EN Koffi, MP Chipperfield, W Winiwarter, and EA Davidson. 2019. ’Acceleration of global N2O emissions seen from two decades of atmospheric inversion’, Nature Climate Change, 9: 993–98.

Tian, Hanqin, Rongting Xu, Josep G Canadell, Rona L Thompson, Wilfried Winiwarter, Parvadha Suntharalingam, Eric A Davidson, Philippe Ciais, Robert B Jackson, and Greet Janssens-Maenhout. 2020. ’A comprehensive quantification of global nitrous oxide sources and sinks’, Nature, 586: 248–56.

Tomlinson, Kira L, Tania Wong Fok Lung, Felix Dach, Medini K Annavajhala, Stanislaw J Gabryszewski, Ryan A Groves, Marija Drikic, Nancy J Francoeur, Shwetha H Sridhar, and Melissa L Smith. 2021. ’Staphylococcus aureus induces an itaconate-dominated immunometabolic response that drives biofilm formation’, Nature communications, 12: 1–13.

Uritskiy, Gherman V, Jocelyne DiRuggiero, and James Taylor. 2018. ’MetaWRAP—a flexible pipeline for genome-resolved metagenomic data analysis’, Microbiome, 6: 1–13.

Van Elsas, Jan Dirk, Mario Chiurazzi, Cyrus A Mallon, Dana Elhottovā, Václav Krištůfek, and Joana Falcão Salles. 2012. ’Microbial diversity determines the invasion of soil by a bacterial pathogen’, Proceedings of the National Academy of Sciences, 109: 1159–64.

Veith, Paul D, Michelle D Glew, Dhana G Gorasia, and Eric C Reynolds. 2017. ’Type IX secretion: the generation of bacterial cell surface coatings involved in virulence, gliding motility and the degradation of complex biopolymers’, Molecular Microbiology, 106: 35–53.

Winiwarter, Wilfried, Lena Höglund-Isaksson, Zbigniew Klimont, Wolfgang Schöpp, and Markus Amann. 2018. ’Technical opportunities to reduce global anthropogenic emissions of nitrous oxide’, Environmental Research Letters, 13: 014011.

Wu, Yu-Wei, Blake A Simmons, and Steven W Singer. 2016. ’MaxBin 2.0: an automated binning algorithm to recover genomes from multiple metagenomic datasets’, Bioinformatics, 32: 605–07.

Wuebbles, Donald J. 2009. ’Nitrous oxide: no laughing matter’, Science, 326: 56–57.

Yoon, Seok-Hwan, Sung-min Ha, Jeongmin Lim, Soonjae Kwon, and Jongsik Chun. 2017. ’A large-scale evaluation of algorithms to calculate average nucleotide identity’, Antonie Van Leeuwenhoek, 110: 1281–86.

Yu, Fengchao, Sarah E Haynes, and Alexey I Nesvizhskii. 2021. ’IonQuant enables accurate and sensitive label-free quantification with FDR-controlled match-between-runs’, Molecular & Cellular Proteomics, 20.

