## Supplementary material 2 for "Meta-omics analyses of dual substrate enrichment culturing of nitrous oxide respiring bacteria suggest that attachment and complex polysaccharide utilisation contributed to the ability of *Cloacibacterium* strains to reach dominance"

**Supplementary material 2**. NO concentrations in liquid during enrichments.

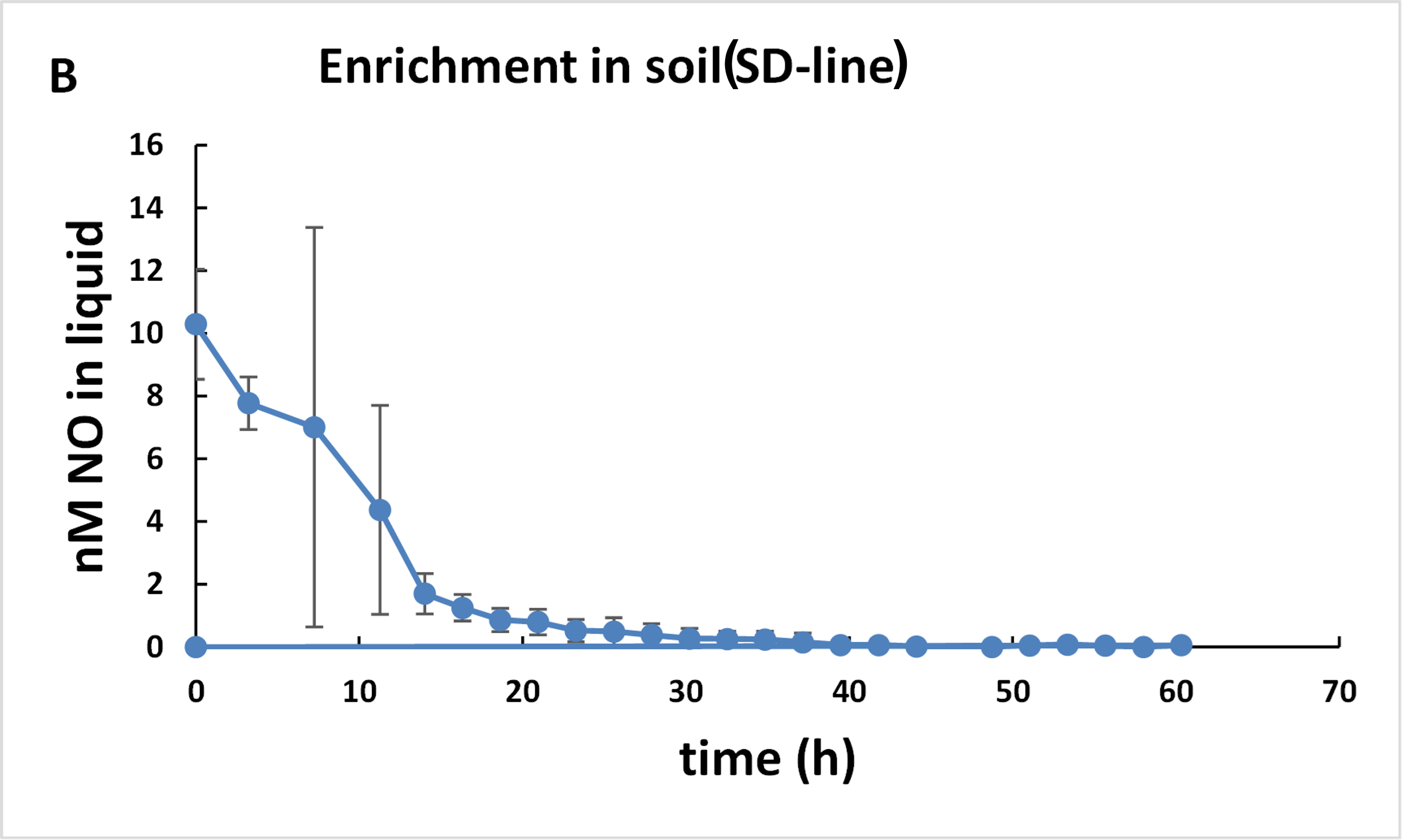

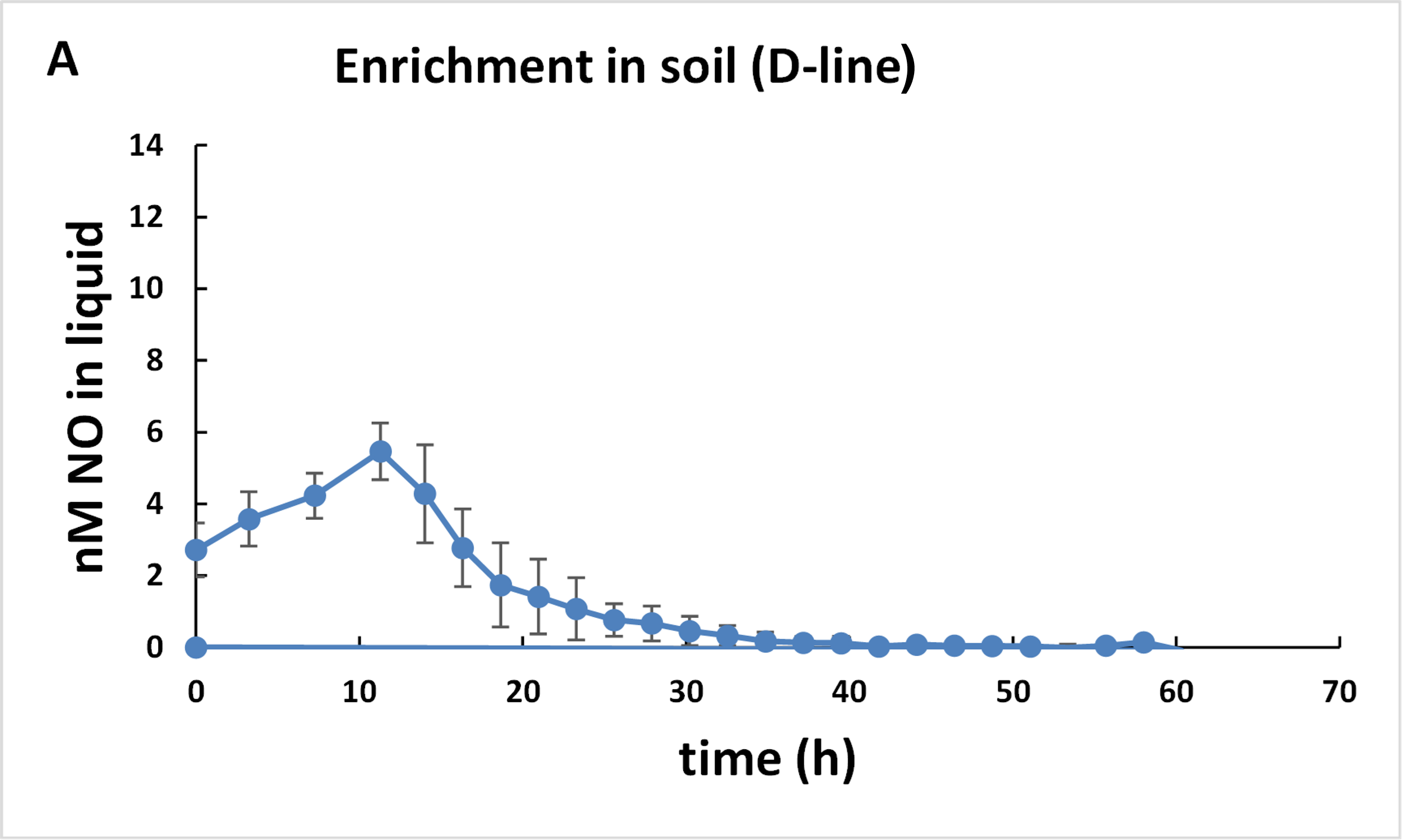

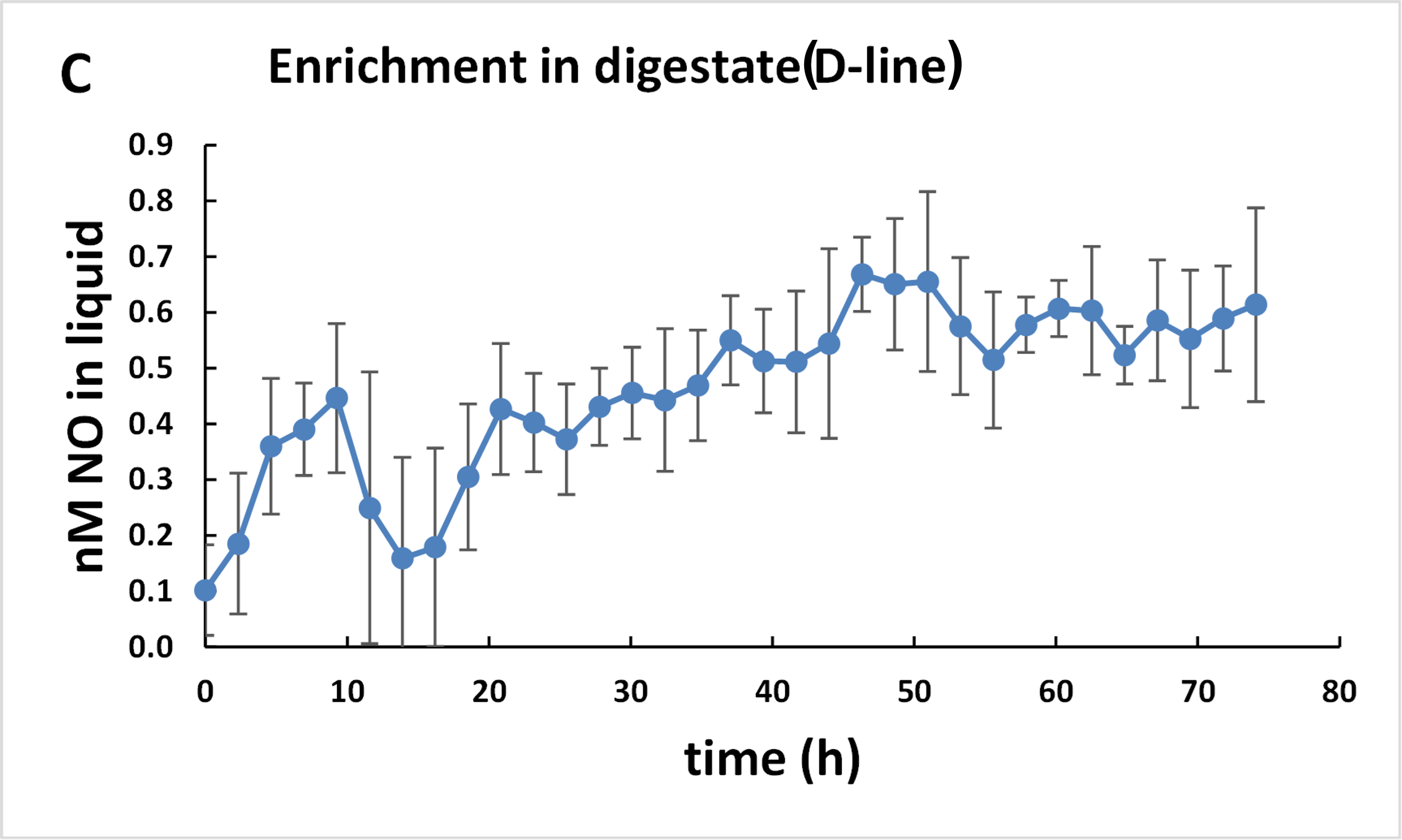

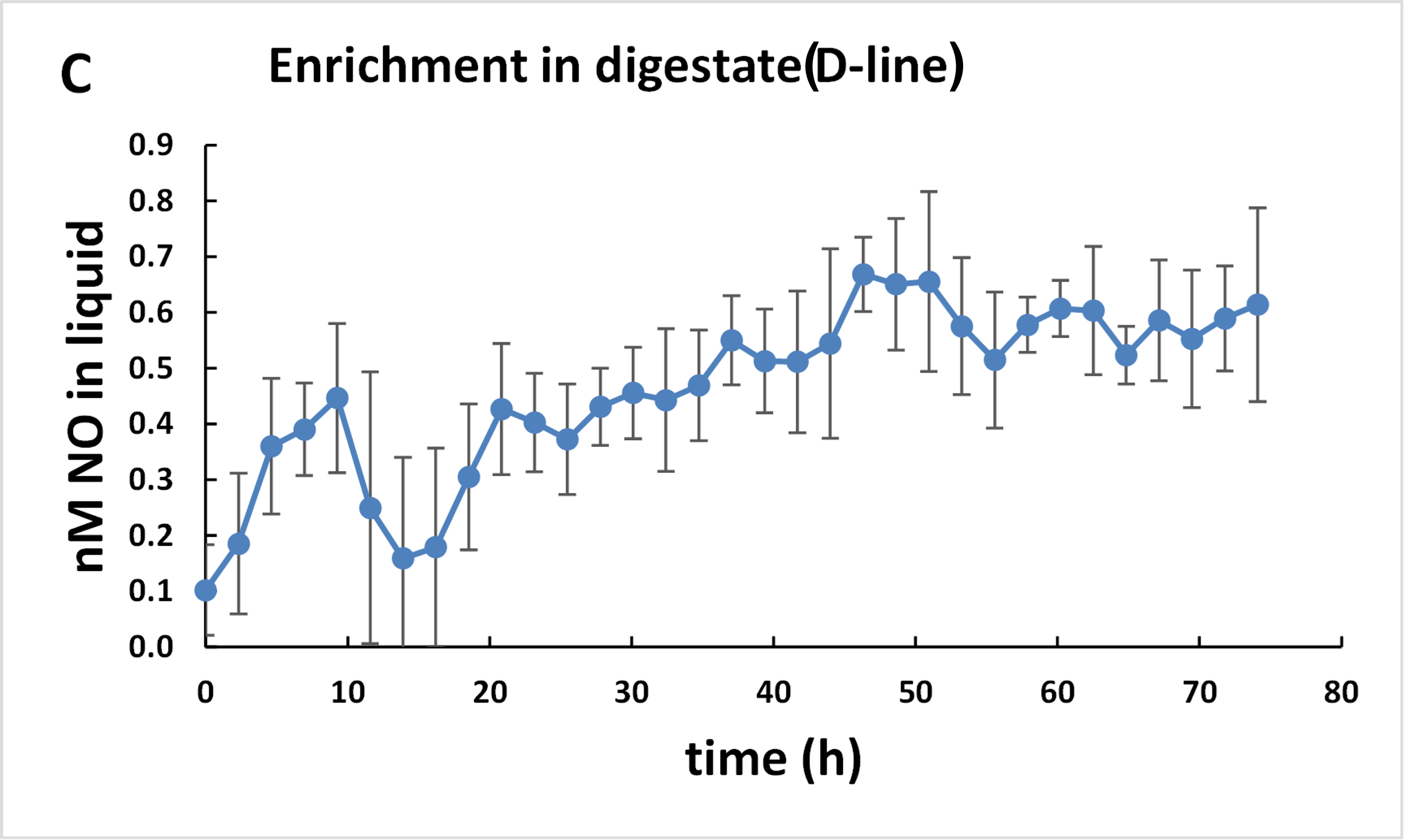

**Measured NO during the enrichment culturing.** The panels show NO-concentrations in the liquid plotted against time (average of 7 replicate vials for each panel, with standard deviation indicated by error bars). Panels A and B show the results for the first enrichment in soil (cycle 1). Panels C and D show the results for the subsequent enrichment in digestate. The subsequent enrichments in soil and digestate gave nearly identical results (enrichments in soil: NO 3-6 nM initially, declining to zero after 10-15 h; enrichments in digestate: marginal NO initially, but increasing gradually to 0.5-0.7 nM. During these enrichments, O_2_ was present initially (2 vol% in headspace) but declined to near zero after 10-15 hours. Oxygen leakage by sampling was 80 nmol O_2_ vial^-1^ for each gas sampling.
